# Immediate and Long-Term Effects of Tibial Nerve Stimulation on the Sexual Behavior of Female Rats

**DOI:** 10.1101/2022.06.20.496650

**Authors:** Lauren L. Zimmerman, Georgios Mentzelopoulos, Hannah Parrish, Vlad I. Marcu, Brandon D. Luma, Jill B. Becker, Tim M. Bruns

**Affiliations:** Department of Biomedical Engineering, University of Michigan, Ann Arbor, MI, USA; Biointerfaces Institute, University of Michigan, Ann Arbor, MI, USA; Department of Electrical Engineering and Computer Science, University of Michigan, Ann Arbor, MI, USA; Department of Engineering Physics, University of Michigan, Ann Arbor, MI, USA; Michigan Neuroscience Institute, University of Michigan, Ann Arbor, MI, USA; Department of Psychology, University of Michigan, Ann Arbor, MI, USA

**Author notes:** **Authorship Statements:** Dr. Zimmerman, Dr. Becker, and Dr. Bruns designed the study. Dr. Zimmerman and Dr. Bruns obtained funding for the study. Dr. Zimmerman, Mr. Mentzelopoulos, Ms. Parrish, Mr. Marcu, and Mr. Luma performed the experimental sessions. Dr. Zimmerman, Mr. Mentzelopoulos, Ms. Parrish, and Mr. Marcu analyzed the data. Dr. Zimmerman and Dr. Bruns drafted the manuscript. All authors reviewed and approved the final manuscript. **Corresponding Author:** Tim M. Bruns, 2800 Plymouth Rd, NCRC B10-A169, Ann Arbor, MI 48109-2800.

**Keywords:** Electric Stimulation, Sexual Behavior, Animal, Tibial Nerve, Sexual Dysfunction, Physiological, Rats

## Abstract

**Objectives:** There are limited treatment options for female sexual dysfunction (FSD). Percutaneous tibial nerve stimulation (PTNS) has shown improvements in FSD symptoms in neuromodulation clinical studies, but the direct effects on sexual function are not understood. This study evaluated the immediate and long-term effects of PTNS on sexual motivation and receptivity in a rat model of menopausal women. Our primary hypothesis was that long-term PTNS would yield greater changes in sexual behavior than short-term stimulation.

**Materials and Methods:** In two Experiments, after receiving treatment, we placed ovariectomized female rats in an operant chamber in which the female controls access to a male by nose poking. We used five treatment conditions, which were with or without PTNS and no, partial, or full hormone priming. In Experiment 1, we rotated rats through each condition twice with behavioral testing immediately following treatment for 10 weeks. In Experiment 2, we committed rats to one condition for 6 weeks and tracked sexual behavior over time. We quantified sexual motivation and sexual receptivity with standard measures.

**Results:** No primary comparisons were significant in this study. In Experiment 1, we observed increased sexual motivation but not receptivity immediately following PTNS with partial hormone priming, as compared to priming without PTNS. In Experiment 2, we observed trends of increased sexual receptivity and some sexual motivation metrics when PTNS was applied long-term with partial hormone priming, as compared to hormone-primed rats without stimulation.

**Conclusions:** PTNS combined with hormone priming shows potential for increasing sexual motivation in the short-term and sexual receptivity in the long-term in rats. Further studies are needed to examine variability in rat behavior and to investigate PTNS as a treatment for FSD in menopausal women.

## Introduction

Female sexual dysfunction (FSD) affects a significant number of women. Around 40% of women present one or more sexual dysfunction symptoms, with low sexual desire and arousal being the most common complaints.^1–4^ FSD is more common in perimenopausal and postmenopausal women, at a rate of around 50%.^4^ FSD can have a serious impact on women’s quality of life.^3^ Despite a significant patient population, there are limited treatment options available.^5^ For several years, neuromodulation techniques that apply electrical stimulation to nerves to treat conditions have shown evidence of being able to treat female sexual dysfunction.^6^ However, little research has been done on how best to utilize this treatment or to examine its mechanisms.

Percutaneous tibial nerve stimulation (PTNS) is a minimally invasive neuromodulation therapy that has been utilized as a treatment for lower urinary tract dysfunctions such as overactive bladder for several decades.^7^ In this therapy, electrical stimulation is delivered to the tibial nerve near the ankle with a percutaneous wire. PTNS is typically delivered in 30-minute stimulation sessions once a week for 12 weeks with periodic maintenance sessions thereafter. Positive benefits begin to present after several weeks of stimulation.^7^ In some studies of PTNS for bladder dysfunction, a secondary outcome of improving sexual dysfunction symptoms has been observed.^8^ Recently we conducted a pilot clinical study of skin-surface PTNS and dorsal genital nerve stimulation (DGNS) in women with FSD but no bladder dysfunction to determine if improvements in sexual functioning can be achieved without concomitant improvement in bladder function.^9^ In that study, the subjects receiving PTNS had increases in their sexual functioning, most significantly in the genital arousal subdomains of arousal, lubrication, and orgasm.^9^ However, the mechanism of these improvements are not understood.

In a previous preclinical study, we investigated a possible mechanism of action by evaluating the effect of tibial nerve stimulation on the vaginal blood perfusion of anesthetized rats.^10^ We showed that long durations (20-40 minutes) of tibial nerve stimulation at 20 Hz can lead to prolonged increases in vaginal blood perfusion, as seen by laser Doppler flowmetry. These increases in blood flow may explain why improvements were seen in the genital arousal components of women with FSD, though in a recent study we found that 20 minutes of tibial nerve stimulation did not have a direct effect on vulvar blood perfusion of anesthetized rats.^11^ Recently we also showed that twice-weekly 30-minute tibial nerve stimulation sessions can maintain vaginal blood flow for several weeks after removal of the ovaries.^12^ While that study suggested that hormones are needed for a longer-term genital arousal benefit from tibial nerve stimulation, the relative hormone amount needed is unknown. Additionally, there have been no studies evaluating the effect of tibial nerve stimulation on the sexual behavior of rats, which may have implications for women.

Here, we investigate whether tibial nerve stimulation can lead to increases in sexual motivation and receptivity. It has been shown that tactile clitoral and vaginal stimulation can modulate sexual behavior^13,14^, but it is unknown whether tibial nerve stimulation can have a similar effect. As ovariectomized rats have been shown to have reduced vaginal blood flow as well as reduced sexual motivation^15,16^, they are used as our model for sexual dysfunction. This model is a standard preclinical model for post-menopausal women,^17^ who frequently have deficits in sexual function.^3^ This animal model does not represent the entire range of FSD symptoms that may be experienced after menopause. Ovariectomized rats have lower sexual motivation and receptivity than intact rats, but their sexual behavior can be restored through hormone priming.^18^

It is unclear whether tibial nerve stimulation would be most beneficial in treating FSD as an on-demand treatment right before sexual activity or as a long-term treatment that leads to improvements over time, similar to PTNS clinical use for bladder dysfunction. In this study, we performed two experiments: one that evaluated the sexual behavior of rats immediately after receiving percutaneous tibial nerve stimulation to investigate the short-term impact of stimulation, and a second that evaluated the sexual behavior of rats over time with repeated tibial nerve stimulation to evaluate the long-term impact of stimulation. The effects of stimulation were evaluated in animals in separate experimental conditions with and without hormone priming using a previously developed method for quantifying the sexual motivation of female rats.^18^

## Materials and Methods

### Animals and Preparation

In this study we used a total of 28 adult female (200-300 g) and 12 adult male (300-400 g) Sprague Dawley rats (Charles River Laboratories, Wilmington, MA, USA), which were split among Experiments 1 and 2 as described below. Animals were housed in same-sex pairs in standard laboratory cages and maintained on a 14:10 hour light:dark cycle (lights off at 12:00 PM) with free access to chow and water. All female rats were bilaterally ovariectomized under isoflurane anesthesia (1-5%) and given carprofen (5 mg/kg subcutaneous) for analgesia for two days following surgery. Beginning 4 days after surgery, we took vaginal epithelium samples via daily saline lavage for 10 days to confirm a complete ovariectomy. We conducted all procedures in accordance with the National Institutes of Health guidelines on laboratory animal use and care, using a study protocol approved by the University of Michigan Institutional Committee on Use and Care of Animals.

### Behavioral Apparatus and Training

The behavioral testing apparatus used in this study is described in Cummings & Becker 2012.^18^ The apparatus consists of a dual-compartment Plexiglas operant chamber. Separating the compartments is a horizontal sliding door that is controlled by rodent nose poking on the female side. Animals are tracked using an overhead monochrome board camera (DMM 72BUC02-ML, The Imaging Source, Charlotte, NC, USA) connected to a computer running the ANY-maze program (Stoetling Co. Inc., Wood Dale, IL, USA), which tracks the animal, collects nose poke information, and controls the door. The smaller chamber is the female side, where the female is free to roam. There are two nose poke holes on the female side, one active and one inactive. The active hole has a corresponding light cue. Nose poking in the active hole in a manner fitting the operant schedule opens the door, giving the female access to the larger side, which contains a tethered male who cannot reach into the female side. Figure 1 shows a diagram of the behavioral testing apparatus.

**Figure 1.**
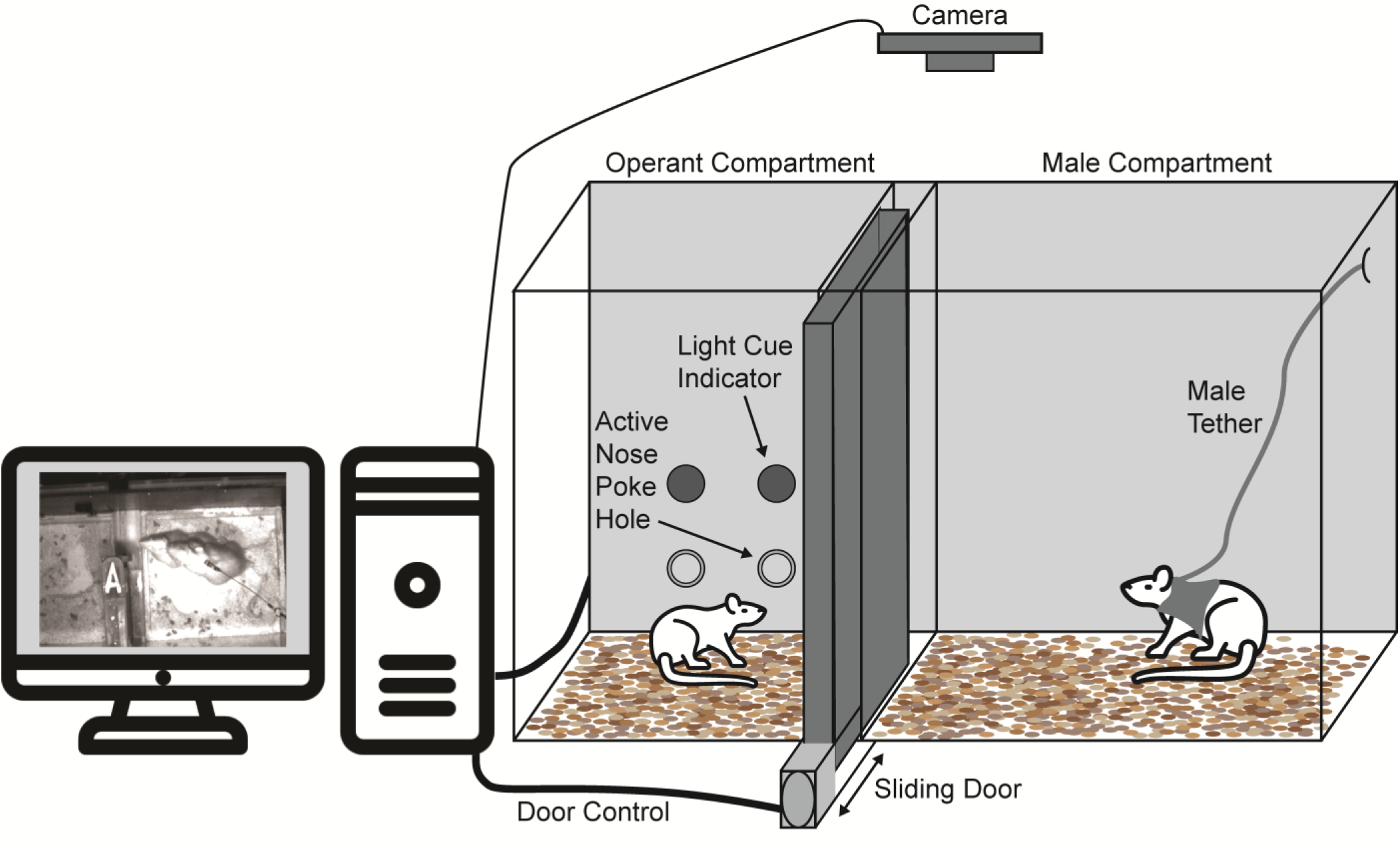
Behavioral testing apparatus, as developed in Cummings & Becker 2012.^18^ Female rats nose poke in an active hole according to an operant schedule to open the sliding door and access the tethered male rat. An overhead camera records activity for analysis by the ANY-maze program on the computer.

We trained all females once a week for 4 weeks before the start of testing with the same hormone priming paradigm to induce consistent pre-testing sexual receptivity. Females were hormone-primed with 10 μg estradiol benzoate (EB) 48 and 24 hours prior and 500 μg progesterone (P) 4-5 hours before each training session. We gave the same doses to all animals, regardless of animal weight, as done in prior hormone-priming studies.^18–22^ We trained females to nose poke in the active hole to open the sliding door on a fixed ratio (FR) and fixed interval (FI) schedule in 30-minute sessions. With each nose poke there is a light cue for 1s. Training females began at FR1, which requires a single nose poke to open the door, and then advanced to FR5, which requires 5 nose pokes to open the door. After the animals were successful on the FR5 schedule they were changed to a fixed interval of 15 seconds (FI15), in which an initial nose poke began a 15-second interval during which nose pokes are recorded.^19^ The door is opened at the conclusion of the FI15 interval if the female exhibited an additional nose poke within 5s after the termination of the 15s interval, otherwise the interval would reset. We used FI15 as the schedule for experimental testing. The operant response (nose poking) schedule with a fixed interval allows for a quantitative measure of motivation (number of nose pokes), as previously validated for assessing female rat sexual motivation under different hormone conditions.^18^

### Testing Conditions

Our conditions involved either stimulation (S+) or no stimulation (S-) with no hormones (H-), partial hormone (H+), or full hormone (H++) dosing. We randomly rotated proven breeder males for each female to prevent mate biases. All testing took place within 1-5 hours after the start of the dark cycle (12 PM).

### Hormones

We administered 10 μg EB in 0.1 mL peanut oil subcutaneously 48 and 24 hours prior to testing in animals receiving hormones (S-H+, S+H+, S-H++), and either 100 μg (partial hormone, H+) or 500 μg (full hormone, H++) P in 0.1 mL peanut oil subcutaneously 4-5 hours prior to testing. It has been shown that the sexual receptivity in female rats can be scaled with P dose.^20^ We chose the 100 μg P (H+) dose to represent approximately half of the maximum receptivity a female can exhibit, and 500 μg (H++) to represent maximum receptivity.^20^ We used the H+ dosing to allow room for increases in sexual receptivity when tibial stimulation is applied. In each testing week, we administered vehicle injections of 0.1 mL peanut oil 48, 24, and 4-5 hours prior to testing control H-animals.

### Stimulation

We delivered electrical stimulation to the right posterior tibial nerve via a bipolar wire electrode (EMG hook electrode, Microprobes for Life Science, Gaithersburg, MD) connected to an Isolated High-Power Stimulator (Model 4100, A-M Systems, Sequim, WA, USA). Each week, we anesthetized every animal (S- and S+) with 5% isoflurane for 10 minutes. In S+ animals we placed the wire percutaneously while under anesthesia in the right hind leg near the tibial nerve, between the gastrocnemius tendon and the tibia, immediately proximal to the ankle.^23^ We wrapped surgical tape around the leg where the wire exited the skin for reinforcement and to prevent the rat from chewing the wire. We gave S-animals the same taping as a control but the wire was not placed. We identified the motor threshold for electrical stimulation which elicited a toe twitch. Once the wire was secured and the threshold was found during the 10 min isoflurane interval, we turned off isoflurane and placed animals in open-top cages to receive stimulation while awake. We delivered stimulation at twice the motor threshold with biphasic, 20 Hz, 200 μs pulse-width stimulation for 30 minutes. We used these stimulation parameters to mimic parameters used in rat studies investigating the effect of tibial nerve stimulation on both sexual^10,12^ and bladder functioning.^24,25^

### Data Analysis

We obtained the number of nose pokes per test, nose pokes per interval, nose-poke latencies (time to first nose poke), and number of attempted and completed FI15s intervals from the ANY-maze software program. Two observers who were blinded to the experimental condition of each rat scored videos using Solomon Coder (version: beta 19.08.02, https://solomon.andraspeter.com). Observers scored the time spent in each chamber area (female area, male area, and in doorway), counts of mounts and intromissions by the males, and lordosis expressions by the females. When lordosis was performed, observers scored it as partial (1-point or 2-point lordosis intensity) or full (3-point lordosis intensity).^21^ We calculated the Lordosis (LQ) as the number of lordosis expressions (partial and full) divided by the number of mounts, as a percentage. We used the average value between the two observers for each measure. We analyzed the data using MATLAB R2018a (Natick, MA, USA).

### Experiment 1

We designed Experiment 1 to test the immediate effect of stimulation on sexual behavior by performing testing immediately following tibial nerve stimulation. We used eight Sprague Dawley females (250-300 g) in this experiment. We used four adult Sprague Dawley males who were proven breeders as stimulus males. Experiment 1 testing took place over 10 weeks. We randomly assigned each female rat to undergo each of the five conditions twice across the study duration (Figure 2). S+ animals received one round of stimulation in a testing week. We delivered stimulation from 35 minutes prior to behavioral testing until 5 minutes prior to testing to allow time to remove the wire and tape and transport female rats to the testing chamber. We held testing days for each female rat 7 days after the previous week’s date. As such we did not expect a high P dose (H++) in one week to have a carry-over effect on a control session without hormones (H-) in the following week. Similarly, we did not expect a condition with electrical stimulation to have a carry-over effect on the condition of the following week, which was electrical stimulation in some cases. We present the data as the average and standard deviation in tables and average and standard error in figures for each condition across all animals. To determine whether the test conditions were related to behavioral metrics, we performed linear mixed effects models using R (Vienna, Austria). We used the treatment group and week as fixed effects and the female rat identifications as a random effect in the model for each behavioral metric. Male rat identifications had no effect on the linear models. We made comparisons between each condition group for each behavioral metric with significance set at p < 0.05.

**Figure 2.**
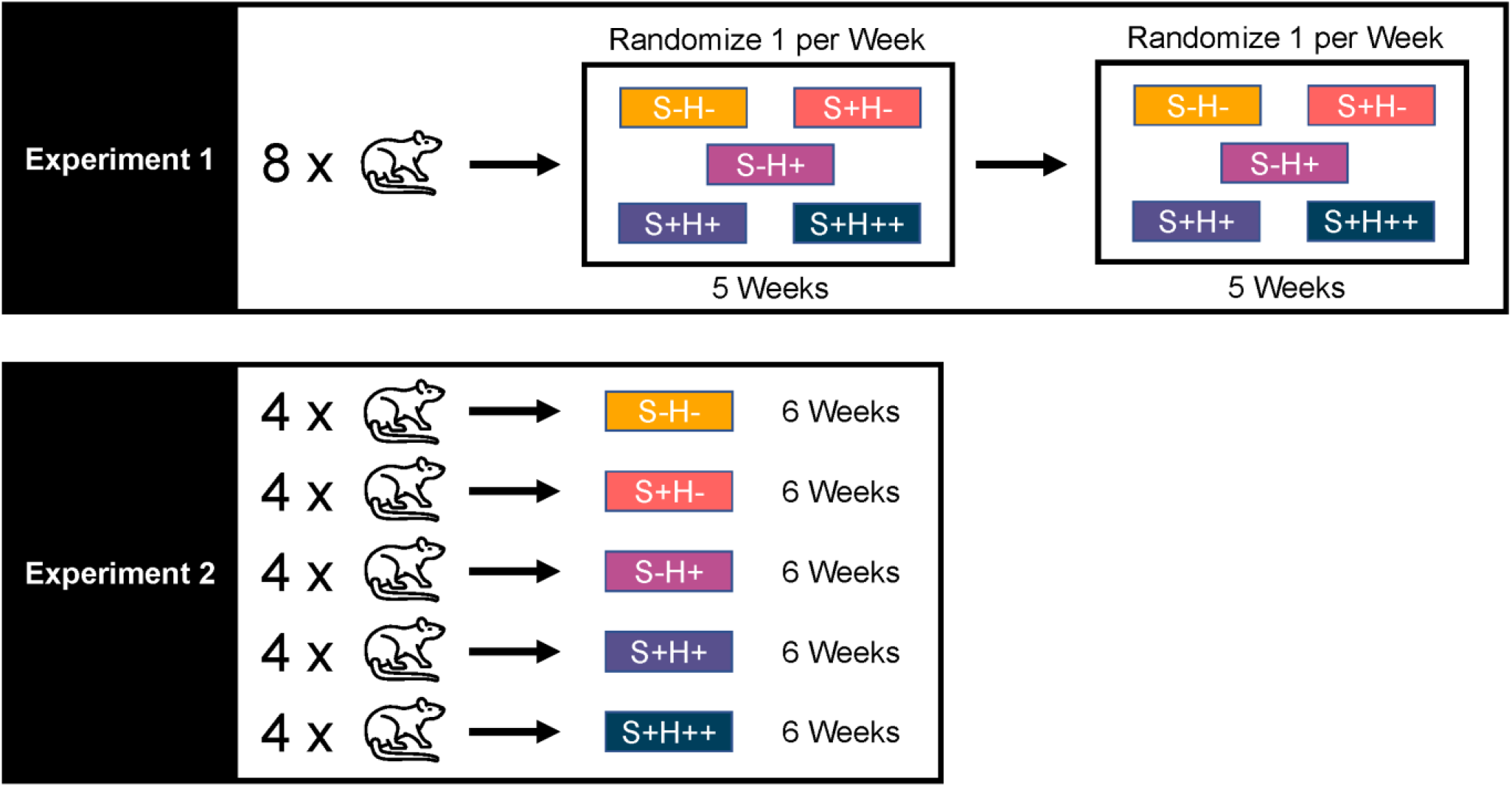
In Experiment 1, rats underwent each of the five conditions in random order in two blocks across the 10-week study duration. Hormone priming and electrical stimulation, when used, were applied to coincide with the behavioral testing session. In Experiment 2, rats were randomly assigned to one of the conditions for the 6-week study duration. Electrical stimulation, when used, was given twice a week to coincide with hormone priming.

### Experiment 2

We designed experiment 2 to test the long-term effects of stimulation by administering stimulation twice a week for six weeks for S+ animals. Twenty Sprague Dawley females (250-300 g) were used in this experiment. We used eight adult Sprague Dawley males who were proven breeders as stimulus males. Experiment 2 testing took place over 6 weeks. We randomly assigned four female rats to each of the five conditions for the entire duration of the study (Figure 2). We gave S+ animals two rounds of stimulation in a testing week, at 48 hours and 24 hours prior to testing to synchronize with EB injections. We determined linear correlations for each metric over the time course of the study. To determine statistical significance, we performed analysis of covariance (ANCOVA) tests using R (Vienna, Austria). We made comparisons between each condition group, evaluating changes over time for each behavioral metric with significance set at p < 0.05.

## Results

All videos and data summary spreadsheets can be found online.^26^ The main comparisons of interest between condition groups were between S+H- and S-H-rats and S+H+ and S-H+ rats. Both comparisons evaluate rats receiving the same hormone treatment (either H- or H+) with S+ rats also receiving stimulation. This comparison allows for an examination of the effects of tibial nerve stimulation on the sexual behavior of rats, depending on hormone condition. Across both Experiments, the majority of main comparisons were not significant, however there were trends in some cases. We report both significant and non-significant comparisons, with p-values given for tests that had statistical significance. In both Experiments, stimulation was typically well tolerated. Some rats lifted their leg or attempted to chew the wires in response to stimulation, indicating some level of stimulation perception or discomfort.

### Experiment 1

All eight rats completed each of the ten weeks of testing. The average amplitude of electrical stimulation administered was 1.18 ± 0.66 mA.

#### Sexual Motivation

Table 1 and Figure 3 present data for motivational metrics for Experiment 1. In this experiment, stimulation in combination with hormone priming led to an increasing trend in sexual motivation compared to rats not receiving stimulation, as indicated by increases in nose pokes per test and nose poke frequency and decreases in initial and inter-interval latencies. S+H+ rats had the shortest initial latency (Figure 3b) and shortest inter-interval latency. S+H+ rats on average had more nose pokes per test (Figure 3a) than S-H+ rats. S+H+ rats had the highest nose poke frequency. Stimulation without hormone priming showed a decreasing trend in sexual motivation compared to hormone primed rats that did not receive stimulation. S+H-rats had significantly fewer nose pokes per interval than S-H+ (p = 0.04) and S-H++ rats (p = 0.02).

**Figure 3.**
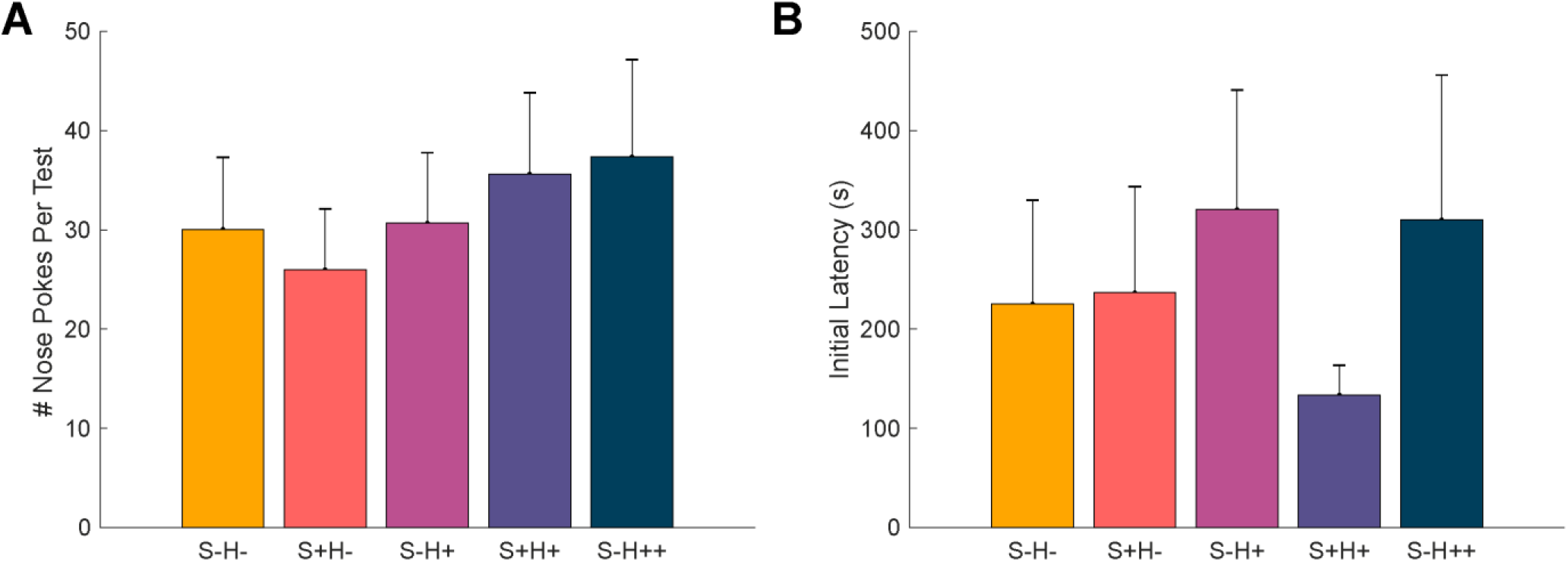
A: Average total nose pokes per test. B: Latency to first nose poke in test session. Error bars represent standard error. There were no significant differences among groups.

**Table 1.** Sexual motivation metrics presented as average ± standard deviation for each treatment group.

|  | **S-H-** | **S+H-** | **S-H+** | **S+H+** | **S-H++** |
| --- | --- | --- | --- | --- | --- |
| **Nose Pokes per Test** | 30.06 | 26.00 | 30.69 | 35.63 | 37.38 |
| | $\pm 28.95$ | $\pm 24.52$ | $\pm 28.33$ | $\pm 32.75$ | $\pm 39.05$ |
| **Nose Pokes per Interval** | 3.73 | **3.20\*** | 3.88 | 3.88 | 4.12 |
| | $\pm 2.54$ | **$\pm 2.17$** | $\pm 2.97$ | $\pm 3.38$ | $\pm 3.65$ |
| **Nose Poke Frequency** | 1.59 | 1.38 | 1.64 | 2.12 | 1.60 |
| | $\pm 1.75$ | $\pm 1.76$ | $\pm 1.52$ | $\pm 2.02$ | $\pm 1.86$ |
| **Initial Latency** | 225.34 | 236.81 | 320.63 | 133.74 | 310.23 |
| | $\pm 418.60$ | $\pm 426.68$ | $\pm 481.90$ | $\pm 120.19$ | $\pm 583.03$ |
| **Inter-Interval Latency** | 98.17 | 99.93 | 118.39 | 89.41 | 95.38 |
| | $\pm 217.47$ | $\pm 170.68$ | $\pm 278.92$ | $\pm 157.65$ | $\pm 194.38$ |
| **Completed Intervals** | 2.19 | 1.81 | 3.06 | 2.81 | 3.25 |
| | $\pm 2.71$ | $\pm 2.29$ | $\pm 3.55$ | $\pm 3.12$ | $\pm 3.97$ |
| **Failed Intervals** | 4.31 | 5.25 | 3.94 | 4.69 | 4.50 |
| | $\pm 3.94$ | $\pm 3.51$ | $\pm 3.70$ | $\pm 3.34$ | $\pm 3.06$ |

Bolded values with \* represent conditions that had significantly different values from other conditions.

#### Sexual Receptivity

Table 2 and Figure 4 show data for receptivity metrics for Experiment 1. Stimulation did not have a positive effect on sexual receptivity, with a decreasing trend in most metrics for hormone primed rats receiving stimulation compared to hormone primed rats not receiving stimulation. S+H+ rats had fewer mounts, intromissions and lower LQs than S-H+ rats and S-H++ rats, as shown in Figure 4a. S+H+ rats on average spent the highest percentage of time with males, as well as highest percentage of time in the doorway (Figure 4b), including significantly more than S-H++ (p < 0.01). When there was no hormone priming, rats receiving stimulation had increasing trends in sexual receptivity metrics compared to rats not receiving stimulation, as S+H-rats had on average more mounts and intromissions than S-H-rats. The most important factor determining sexual receptivity was the level of hormone priming. S-H-had significantly fewer mounts than S-H+ (p = 0.04), S+H+ (p = 0.05) and S-H++ rats (p = 0.01). S-H- and S+H-rats both had significantly fewer intromissions than S-H++ rats (p < 0.01, p = 0.02 respectively).

**Figure 4.**
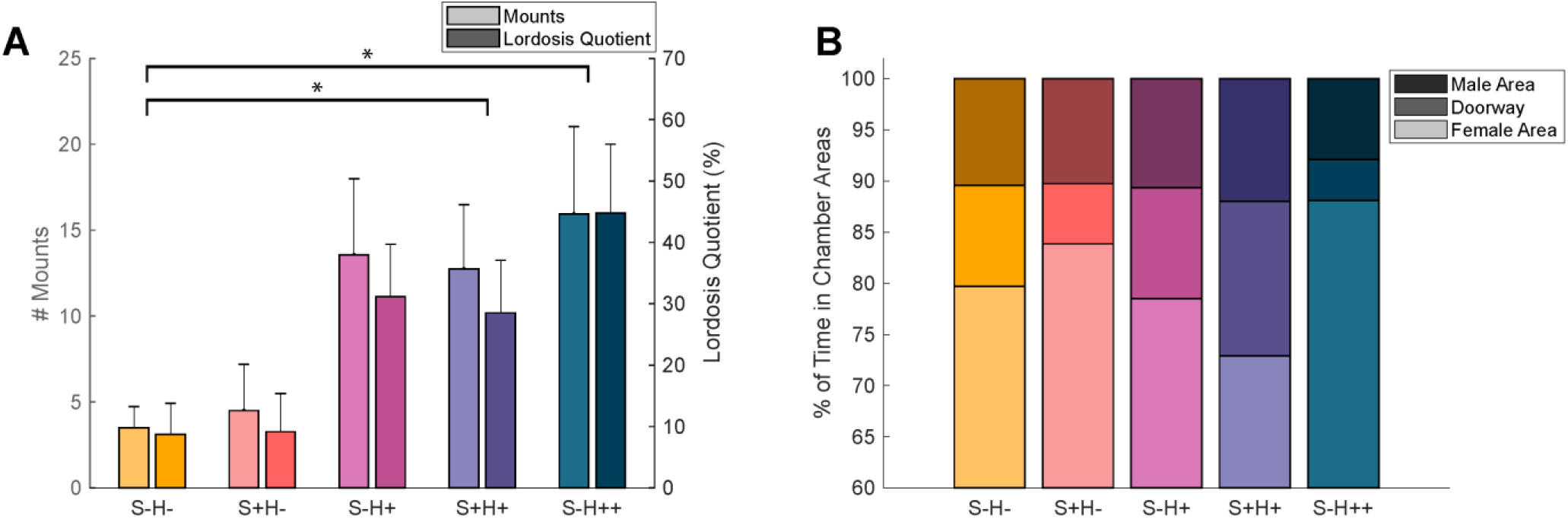
A: Average mounts and lordosis quotient across conditions. Error bars give standard error. Brackets with * represent significant difference between condition groups (p < 0.05) for both mounts and LQ. B: Average percentage of time spent in the male area, doorway, and female area across conditions.

**Table 2.** Average of each receptivity metric across each treatment group in Experiment 1.

|  | **S-H-** | **S+H-** | **S-H+** | **S+H+** | **S-H++** |
| --- | --- | --- | --- | --- | --- |
| **Mounts** | 3.50 | 4.50 | 13.56 | 12.75 | 15.94 |
|  | ± 4.90 | ± 10.81 | ± 17.75 | ± 14.94 | ± 20.35 |
| **Intromissions** | 1.47 | 3.50 | 7.91 | 7.53 | 12.78 |
|  | ± 2.75 | ± 9.64 | ± 11.19 | ± 10.52 | ± 16.88 |
| **Lordosis Quotient (%)** | 8.7 | 9.2 | 31.2 | 28.9 | 44.9 |
|  | ± 20.1 | ± 25.0 | ± 34.1 | ± 35.2 | ± 44.9 |
| **% Time with Male** | 10.43 | 10.26 | 10.66 | 11.96 | 7.87 |
|  | ± 19.24 | ± 18.80 | ± 15.66 | ± 13.57 | ± 10.61 |
| **% Time in Doorway** | 9.85 | 5.87 | 10.81 | 15.14 | 4.00 |
|  | ± 11.03 | ± 6.97 | ± 11.34 | ± 14.87 | ± 5.47 |
| **% Time Alone** | 79.73 | 83.87 | 78.53 | 72.89 | 88.12 |
|  | ± 26.39 | ± 22.38 | ± 24.21 | ± 25.29 | ± 5.47 |

### Experiment 2

All twenty rats completed each of the six weeks of treatment and testing. The average amplitude of stimulation administered was 3.24 ± 2.46 mA. Stimulation amplitude increased over time, with an average amplitude of 2.75 ± 1.07 mA in the first stimulation session in Week 1 increasing to 4.90 ± 2.99 mA in the last session in Week 6. There was high variability across metrics in this Experiment. Therefore, we focused our analysis on trends over time for the condition groups. All data is presented as average ± standard deviation per treatment group per week in Tables S1 and S2 of the Supporting Information.

#### Sexual Motivation

Figure 5 shows key sexual motivation metrics. Table 3 shows linear correlation slope values for Experiment 2. In this Experiment, rats receiving stimulation and hormone priming had increasing trends in sexual motivation over time compared to rats receiving hormone priming but no stimulation. S+H+ rats had the highest slope for nose pokes per interval across testing sessions, and a lower slope than S-H+ for nose pokes per test (Figure 5a) and completed intervals (Figure 5c). S+H+ rats had a higher slope for nose poke frequency than S-H+ and other conditions except S-H++. S+H+ rats had the most negative slope for initial latency (Figure 5b).

**Figure 5.**
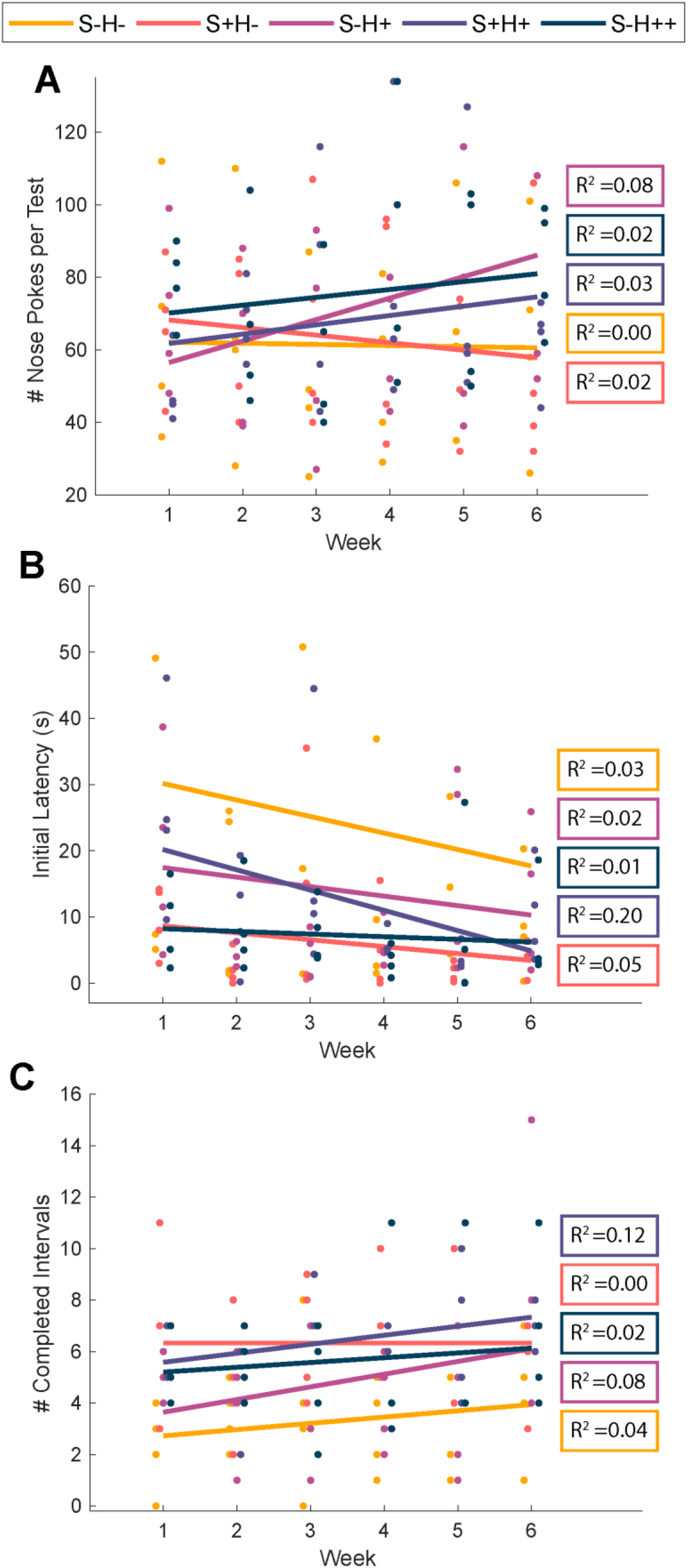
Motivation metrics for Experiment 2 across the 6-week duration. Linear correlation lines across 6 weeks presented with R^2^ values. A: Nose pokes per test. B: Latency to first nose poke of test C: Completed intervals.

**Table 3.** Linear correlation slopes for Experiment 2 sexual motivation and receptivity metrics.

|  | Unit | S-H- | S+H- | S-H+ | S+H+ | S-H++ |
| --- | --- | --- | --- | --- | --- | --- |
| **Motivation Metrics** |  |  |  |  |  |  |
| Nose Pokes per Test | Nose pokes | -0.314 | -2.086 | 5.921 | 2.571 | 2.164 |
| Nose Pokes per Interval | Nose pokes | 0.293 | 0.119 | 0.403 | 0.455 | 0.199 |
| Nose Poke Frequency | Nose pokes/min | 0.294 | -0.015 | 0.463 | 0.661 | 1.357 |
| Initial Latency | Seconds | -2.493 | -1.045 | -1.433 | -3.062 | -0.400 |
| Inter-Interval Latency | Seconds | -1.175 | 1.391 | -0.503 | -1.811 | -1.887 |
| Completed Intervals | Intervals | 0.243 | 0.000 | 0.493 | 0.350 | 0.186 |
| Failed Intervals | Intervals | -1.093 | -0.921 | -0.814 | -1.336 | -0.579 |
| **Receptivity Metrics** |  |  |  |  |  |  |
| Mounts | Mounts | -1.232 | -2.200 | 0.718 | 2.650 | -0.518 |
| Lordosis Quotient | # Lordosis Expressions / # Mounts | -0.078 | -0.062 | -0.083 | 0.024 | -0.060 |
| Intromissions | Intromission s | -0.364 | -0.654 | 0.714 | 2.425 | -1.296 |
| % Time with Male | % of 30-minute test | 0.366 | -2.084 | -0.841 | 1.148 | -0.649 |
| % Time in<br>Doorway | % of 30-minute test | 1.611 | 2.050 | 0.774 | 1.911 | 1.746 |
| % Time Alone | % of 30-minute test | -1.977 | 0.034 | 0.067 | -3.059 | -1.098 |

#### Sexual Receptivity

Figure 6 shows key sexual receptivity metrics. Table 3 shows linear correlation slopes. Rats receiving stimulation and hormone priming had increasing trends in sexual receptivity over time compared to rats receiving hormone priming but no stimulation. S+H+ rats had the highest positive slope for mounts (Figure 6a), intromissions (Figure 6b), lordosis quotient (Figure 6c), and time spent with male (Figure 6e), as well as the most negative slope for time alone (Figure 6d).

**Figure 6.**
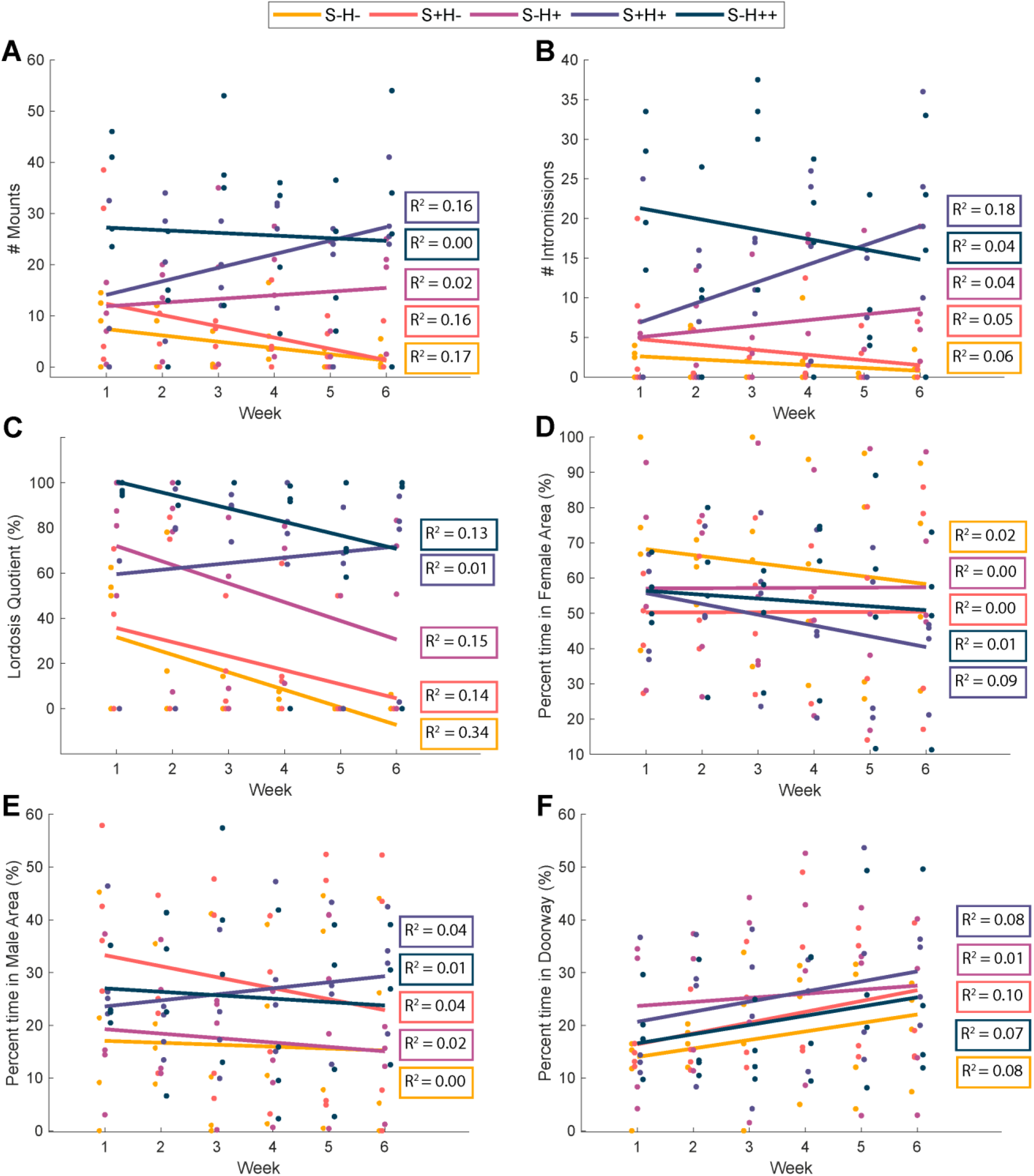
Sexual receptivity metrics for Experiment 2. Linear correlation across 6 weeks. R^2^ values presented. A. Mounts. B. Total lordosis quotient. C. Intromissions. D. Percentage time alone. E. Percentage of time with male. F. Percentage of time in doorway.

## Discussion

This study was the first to investigate tibial neuromodulation in a preclinical sexual motivation and sexual behavior paradigm. To our knowledge, this study was also the first study to evaluate the combination of neuromodulation and hormone dosing as a potential sexual dysfunction treatment in either a clinical or preclinical setting. In Experiment 1, we found trends of increasing sexual motivation immediately following tibial nerve stimulation, but not receptivity. In Experiment 2, we found stronger trends of increasing sexual receptivity when tibial nerve stimulation was delivered long-term on a twice weekly basis. In both Experiments, most comparisons between conditions were not statistically significant. Thus, while we observed average differences or trends between main comparisons of interest that supported our hypotheses, we cannot confidently state that these observations were not due to random variations in animal behavior.

Regarding the potential immediate effects of stimulation on sexual motivation, as seen by S+H+ rats having the shortest initial latency (Figure 3b), shortest inter-interval latency, and the highest nose poke frequency compared to other treatment groups, and more nose pokes per test than S-H+ rats (Figure 3a), our results suggest an increase in motivation to reach the male immediately after receiving stimulation. When stimulation was delivered long-term in Experiment 2, S+H+ rats had higher increases in some of the motivational metrics over time (decreased initial latency (Figure 5b), increased nose pokes per interval) but did not outperform on other metrics (nose pokes per test (Figure 5a), completed intervals (Figure 5c)) compared to S-H+ and S-H++ rats. While non-significant, these trends suggest a stronger increase in motivation immediately following stimulation compared to an effect over time with long-term stimulation.

No increases in sexual receptivity were seen immediately after stimulation in Experiment 1. S+H+ rats had a similar number of mounts and lordosis expressions as S-H+ rats. Lordosis has been shown to depend on oestrogens followed by progestins^27^, so the lack of immediate impact of tibial stimulation on sexual receptivity is unsurprising. However, strong trends of increasing receptivity were seen when stimulation was applied long term in Experiment 2. S+H+ rats had the highest increases in mounts, lordosis quotient, intromissions, time with male, and decreasing time alone across the five conditions (Figure 6). Further studies repeating this paradigm with larger sample sizes or clinical studies comparing on-demand versus long-term tibial nerve stimulation are needed to confirm these results. These experiments suggest that long-term PTNS over several weeks may be a better treatment option for women with FSD than an on-demand stimulation treatment paradigm. If cumulative stimulation is necessary for increases in sexually receptive behavior, as seen in Experiment 2, it is possible that rotating the rats though the conditions in Experiment 1 was unpriming the rats for each testing session.

As the trends described above are only seen in S+H+ and not S+H-rats, our results suggest that tibial nerve electrical stimulation alone does not alter sexual motivation or receptivity. Rather, a combination of stimulation and hormone priming may be an effective treatment for FSD. PTNS combined with partial hormone dosing may reduce the need for hormone treatment at full dosing levels. S+H-exhibited the least sexual motivation and receptivity out of all treatment groups for some metrics (nose pokes per test, nose pokes per interval, completed intervals). These results may indicate that stimulation alone may have an adverse effect on behavior, particularly immediately following stimulation. It is possible that the wire placement and electrical stimulation were disorienting if it was not combined with some hormones to motivate sexual behavior.

There are several possible explanations for these observations. Diminished genital blood flow is a common physiological factor in FSD^28^ and is a side effect of ovariectomizing female rats.^16^ Hormone priming has been shown to maintain vaginal and pelvic floor muscle health after ovariectomy,^29,30^ which should have yielded consistent muscle health across H+/H++ animal groups in Experiment 2 but may have contributed to variability in Experiment 1 animals. Tibial nerve stimulation has been shown to increase vaginal blood flow^10^ but not vulvar blood flow^11^ in anesthetized rats. Stimulation delivered in sessions repeated across a long-term interval could be continually improving genital blood flow, leading to more positive sexual experiences for female rats. Recently we showed that tibial nerve stimulation after ovariectomy can maintain vaginal blood flow increases for several weeks but not to six weeks^12^, aligning with findings of this study that hormone priming may also be needed for long-term benefits. Electrical stimulation may strengthen the spinal circuit for sexual function or may lead to cortical changes. Cortically, the electrical stimulation could be leading to local synthesis of estradiol, though our recent study did not find changes in serum estradiol levels^12^, or activation of estrogen-concentrating regions such as the medial preoptic area, ventromedial hypothalamus, or the medial amygdala that are typically activated during sexual stimulation.^31^ The increases in lordosis expressions are more likely to be driven by cortical changes than increases in genital blood flow, as lordosis is a cortically-controlled behavior.

This study had several limitations. In both experiments, there was high variability in the measurements. Some rats were occasionally unresponsive or had difficulty opening the door following the FI15 interval. Rats in these instances would nose poke several times in a clear attempt to open the door but would miss the 5s latency period after the 15s interval in which a nose poke was required to open the door. When rats consistently failed in this manner, there were no interactions with the male or changes in apparatus areas. This lowered their metrics for sexual receptivity despite showing motivation. This issue could be related to inadequate training and could be remedied by a higher number of training sessions prior to testing. Although S-H++ rats generally performed better than S-H+ rats in Experiment 1 metrics, we did not expect to see S-H+ rats perform better over time in Experiment 2 for most metrics. This contrast may also be due to the variability in measures as the two conditions were not significantly different. Future studies could use a simpler chamber to reduce variance.^22,32^ We did not find statistical significance in many of the treatment groups, due in part to unresponsive sessions. Larger sample sizes may have led to clearer outcomes in the measures.

Another limitation of this study is that verification of consistent stimulation for an entire session was challenging with awake rats, as when the rats were ambulatory the stimulation-driven toe twitches in their hind limbs became imperceptible. The percutaneous wires were loosely anchored and had the possibility of shifting once inserted. We limited the current amplitude to twice the motor threshold, to help reduce the potential for adverse behavioral responses. Studies with anesthetized rats often use higher stimulation amplitudes.^10,24,25^ These higher amplitudes may have more consistent recruitment of the full tibial nerve but may cause greater discomfort for awake animals. It is possible that inconsistent tibial nerve stimulation occurred across the study due to wire positioning or partial recruitment of relevant pathways in the tibial nerve. As this was the first study to test the effects of long-term tibial nerve stimulation on sexual behavior, there was not precedent for the duration of treatment. It is possible that further benefits would be seen if the stimulation was delivered for longer intervals or more sessions than the 6-week treatment period used here.

The average stimulation amplitude was higher in Experiment 2 than Experiment 1, and the average amplitude increased across the duration of Experiment 2. Rats who received stimulation in Experiment 2 were being stimulated more frequently and more regularly than rats in Experiment 1. This increase in stimulation sessions could have led to an increased inflammatory response or scar tissue buildup that necessitated a higher amplitude to achieve motor contractions due to higher tissue resistance.

Future research to replicate or extend this study with a higher number of animals could lead to more conclusive results. Inclusion of intact, cycling rats would allow for a direct comparison to normal behavior across the estrus cycle.^33^ To ensure more consistent stimulation, chronic nerve cuffs could be surgically implanted on the tibial nerve.^34^ Additionally, related studies could examine the effect of tibial nerve stimulation on circulating hormone levels to determine if stimulation is leading to an increase in progesterone synthesis. An increase in hormone production in response to nerve stimulation at other locations has been documented^33,35^ and could explain the increase in sexually receptive behavior over time that we observed here. The vaginal and pelvic floor structural integrity can be examined in a future study ^29,30^ to determine if stimulation has beneficial effects at different hormone levels. The cortical effects of stimulation can also be studied by looking at c-fos activation in the estrogen-containing regions of the brain in response to tibial stimulation.^31^

## Conclusion

In this study we did not observe significant effects in most of the primary comparisons between conditions with and without electrical stimulation of the tibial nerve. We observed trends of increasing sexual motivation but not receptivity immediately after stimulation combined with partial hormone priming. When tibial nerve stimulation was combined with partial hormone priming across weeks, we observed increasing trends in both motivation and receptivity. These trends suggest that long-term tibial nerve stimulation could be a useful treatment modality for treating female sexual dysfunction, especially in post-menopausal populations receiving hormone replacement therapy. Further studies are needed to confirm that these trends are not due to variances in animal behavior during a challenging operant condition.

## Acknowledgements

We thank Dr. Katie Yoest, Sara Bender-Bier, and Alex Ramer for their assistance in setting up this study. This study was funded in part by a fellowship from the Eunice Kennedy Shriver National Institute of Child Health & Human Development of the National Institutes of Health (F31HD094480).

