## supporting information for "Immediate and Long-Term Effects of Tibial Nerve Stimulation on the Sexual Behavior of Female Rats"

**Table S1**. Motivation metrics from Experiment 2 presented as average ± standard deviation per treatment group per week. %Δ W1:W6 represents the percent change from the average in week 1 to week 6 for that metric per condition.

| | Condition | Week 1 | Week 2 | Week 3 | Week 4 | Week 5 | Week 6 | %Δ W1:W6 |
|---|---|---|---|---|---|---|---|---|
| **Nose Pokes per Test** | S-H- | 67.5 ± 33.16 | 65.25 ± 33.78 | 51.25 ± 25.98 | 53.25 ± 23.30 | 66.75 ± 29.35 | 64.00 ± 31.08 | -5.2% |
| | S+H- | 66.5 ± 18.21 | 64.00 ± 22.38 | 67.25 ± 30.21 | 67.25 ± 32.37 | 56.75 ± 20.02 | 56.25 ± 33.81 | -15.4% |
| | S-H+ | 70.25 ± 22.14 | 59.25 ± 23.96 | 60.75 ± 29.78 | 62.25 ± 17.59 | 70.75 ± 34.92 | 104.50 ± 67.75 | 48.8% |
| | S+H+ | 49.00 ± 10.23 | 67.75 ± 10.75 | 76.00 ± 32.95 | 79.50 ± 37.55 | 74.50 ± 35.27 | 62.25 ± 12.63 | 27.0% |
| | S-H++ | 78.75 ± 11.18 | 67.50 ± 25.85 | 59.75 ± 22.29 | 87.75 ± 37.03 | 76.75 ± 28.65 | 82.75± 17.37 | 5.1% |
| **Nose Pokes per Interval** | S-H- | 4.82 ± 3.25 | 3.84 ± 2.90 | 3.80 ± 3.12 | 4.18 ± 4.36 | 5.68 ± 5.98 | 5.69 ± 5.08 | 18.1% |
| | S+H- | 3.86 ± 2.14 | 3.94 ± 2.19 | 4.27 ± 2.34 | 4.56 ± 2.93 | 4.45± 2.04 | 4.33± 2.46 | 12.2% |
| | S-H+ | 3.47 ± 2.07 | 3.95 ± 3.14 | 3.63 ± 2.99 | 3.95 ± 3.32 | 5.05 ± 5.3 | 5.57 ± 4.82 | 60.5% |
| | S+H+ | 3.27 ± 1.77 | 3.43 ± 2.26 | 4.00 ± 2.87 | 5.8 ± 4.07 | 4.73 ± 3.28 | 5.30 ± 2.80 | 62.1% |
| | S-H++ | 4.70 ± 3.13 | 4.03 ± 3.56 | 3.85 ± 2.44 | 5.32 ± 4.48 | 4.80 ± 5.33 | 5.34 ± 6.40 | 13.6% |
| **Nose Poke Frequency** | S-H- | 3.99 ± 3.54 | 3.54 ± 2.29 | 2.93 ± 1.83 | 3.37 ± 1.73 | 5.71 ± 4.64 | 4.66 ± 3.40 | 16.8% |
| | S+H- | 4.87 ± 0.63 | 3.89 ± 1.54 | 4.34 ± 1.33 | 4.67 ± 1.85 | 4.86 ± 1.95 | 4.11 ± 1.68 | -15.6% |
| | S-H+ | 4.70 ± 3.16 | 4.72 ± 4.12 | 4.18 ± 2.64 | 5.42 ± 4.83 | 8.44 ± 9.72 | 5.46 ± 3.3 | 16.2% |
| | S+H+ | 3.22 ± 0.66 | 4.13 ± 1.16 | 4.95 ± 1.99 | 7.09 ± 4.35 | 6.52 ± 2.47 | 5.98 ± 3.55 | 85.7% |
| | S-H++ | 4.60 ± 1.03 | 4.35 ± 1.85 | 4.04 ± 1.01 | 6.06 ± 4.8 | 9.79 ± 12.41 | 10.43 ± 12.25 | 126.7% |
| **Initial Latency** | S-H- | 36.35 ± 37.54 | 13.42 ± 13.61 | 33.53 ± 29.21 | 12.65 ± 16.56 | 36.80 ± 43.31 | 9.05 ± 8.32 | -75.1% |
| | S+H- | 9.73 ± 5.29 | 2.18 ± 2.62 | 13.13 ± 16.34 | 5.28 ± 7.17 | 1.65 ± 1.47 | 4.30 ± 2.81 | -55.8% |
| | S-H+ | 19.50 ± 15.05 | 22.50 ± 36.50 | 4.10 ± 3.78 | 5.73 ± 3.46 | 17.35 ± 15.24 | 12.23 ± 11.10 | -37.3% |
| | S+H+ | 25.88 ± 15.09 | 10.15 ± 8.13 | 17.95 ± 18.03 | 6.98 ± 2.35 | 3.80 ± 1.96 | 10.45 ± 7.28 | -59.6% |
| | S-H++ | 8.90 ± 6.42 | 8.30 ± 7.11 | 7.55 ± 4.66 | 3.40 ± 2.22 | 8.13 ± 13.00 | 7.03 ± 7.73 | -21.0% |
| **Inter-Interval Latency** | S-H- | 82.67 ± 112.99 | 57.82 ± 94.95 | 81.36 ± 100.29 | 75.69 ± 101.68 | 63.23 ± 114.27 | 72.33 ± 94.15 | -12.5% |
| | S+H- | 46.48 ± 75.25 | 42.57 ± 78.69 | 51.71 ± 100.46 | 47.62 ± 104.73 | 66.81 ± 127.79 | 42.49 ± 82.48 | -8.6% |
| | S-H+ | 39.68 ± 55.99 | 45.35 ± 91.68 | 52.14 ± 114.7 | 44.40 ± 70.02 | 51.04 ± 68.51 | 34.29 ± 41.30 | -13.6% |
| | S+H+ | 56.12 ± 67.79 | 31.42 ± 55.24 | 35.64 ± 70.89 | 53.39 ± 79.60 | 42.19 ± 74.74 | 33.43 ± 39.73 | -40.4% |
| | S-H++ | 51.34 ± 84.79 | 42.26 ± 51.71 | 39.31 ± 56.79 | 42.18 ± 54.52 | 53.45 ± 102.9 | 30.84 ± 51.88 | -39.9% |
| **Completed Intervals** | S-H- | 2.25 ± 1.71 | 3.50 ± 1.29 | 3.75 ± 3.30 | 3.00 ± 1.83 | 3.25 ± 2.06 | 4.25 ± 2.50 | 88.9% |
| | S+H- | 7.00 ± 3.27 | 4.75 ± 2.50 | 6.50 ± 2.38 | 7.25 ± 2.06 | 7.00 ± 3.46 | 5.50 ± 1.73 | -21.4% |
| | S-H+ | 5.25 ± 0.96 | 4.00 ± 2.16 | 3.50 ± 2.52 | 4.00 ± 1.83 | 3.75 ± 2.75 | 8.75 ± 4.57 | 66.7% |
| | S+H+ | 5.50 ± 1.00 | 5.00 ± 2.00 | 8.00 ± 1.15 | 6.25 ± 0.50 | 6.75 ± 2.75 | 7.25 ± 0.96 | 31.8% |

| | | | | | | | | |
|---|---|---|---|---|---|---|---|---|
| | S-H++ | 5.75 ± 1.50 | 5.50 ± 1.29 | 4.75 ± 2.22 | 5.50 ± 3.70 | 5.75 ± 3.50 | 6.75 ± 3.10 | 17.4% |
| **Failed Intervals** | S-H- | 11.50 ± 3.42 | 12.75 ± 6.99 | 9.25 ± 2.99 | 9.50 ± 3.11 | 8.25 ± 5.74 | 6.50 ± 3.51 | -43.5% |
| | S+H- | 9.75 ± 6.65 | 10.50 ± 4.20 | 9.00 ± 3.56 | 7.00 ± 4.24 | 5.00 ± 2.16 | 7.00 ± 7.53 | -28.2% |
| | S-H+ | 14.75 ± 3.95 | 10.25 ± 4.43 | 12.75 ± 9.60 | 11.50 ± 6.86 | 9.50 ± 5.32 | 9.75 ± 7.54 | -33.9% |
| | S+H+ | 9.00 ± 5.29 | 13.75 ± 6.80 | 10.75 ± 5.91 | 6.75 ± 4.35 | 8.25 ± 6.90 | 3.75 ± 1.50 | -58.3% |
| | S-H++ | 11.00 ± 5.72 | 10.25 ± 6.40 | 9.75 ± 7.18 | 10.50 ± 4.80 | 9.50 ± 8.96 | 7.25 ± 6.29 | -34.1% |

**Table S2**. Receptivity Metrics from Experiment 2 presented as average ± standard deviation per treatment group per week. %Δ W1:W6 represents the percent change from the average in week 1 to week 6 for that metric per condition.

| | Condition | Week 1 | Week 2 | Week 3 | Week 4 | Week 5 | Week 6 | %Δ W1:W6 |
|---|---|---|---|---|---|---|---|---|
| **Mounts** | S-H- | 9.00 ± 6.42 | 5.25 ± 6.18 | 1.88 ± 3.42 | 6.88 ± 6.68 | 0.88 ± 1.75 | 2.00 ± 2.48 | -77.8% |
| | S+H- | 18.75 ± 18.75 | 3.75 ± 4.97 | 5.13 ± 4.01 | 6.13 ± 7.47 | 4.63 ± 4.50 | 2.63 ± 4.31 | -86.0% |
| | S-H+ | 8.63 ± 6.69 | 13.13 ± 8.53 | 18.63 ± 14.12 | 16.13 ± 10.91 | 8.38 ± 11.15 | 17.00 ± 9.94 | 97.0% |
| | S+H+ | 10.00 ± 15.41 | 22.00 ± 12.62 | 19.00 ± 7.13 | 25.50 ± 9.60 | 18.25 ± 12.34 | 29.50 ± 7.80 | 195.0% |
| | S-H++ | 34.38 ± 10.83 | 13.63 ± 10.86 | 34.38 ± 16.91 | 23.88 ± 13.67 | 20.88 ± 13.20 | 28.50 ± 22.35 | -17.1% |
| **Lordosis Quotient (%)** | S-H- | 41.6 ± 28.2 | 23.7 ± 37.1 | 3.6 ± 7.1 | 2.9 ± 3.7 | 0.0 ± 0.0 | 1.6 ± 3.1 | -96.2% |
| | S+H- | 28.1 ± 34.6 | 39.9 ± 46.3 | 17.5 ± 22.8 | 22.7 ± 28.4 | 12.5 ± 25.0 | 1.0 ± 2.1 | -96.3% |
| | S-H+ | 79.6 ± 21.3 | 68.6 ± 41.7 | 38.0 ± 40.3 | 57.4 ± 31.4 | 12.5 ± 25.0 | 49.5 ± 35.6 | -37.8% |
| | S+H+ | 41.3 ± 49.8 | 64.2 ± 43.6 | 86.9 ± 9.1 | 81.1 ± 14.9 | 55.6 ± 38.6 | 65.8 ± 42.3 | 59.1% |
| | S-H++ | 96.6 ± 2.4 | 97.5 ± 5.0 | 100.0 ± 0.0 | 70.8 ± 47.3 | 74.6 ± 17.8 | 100.0 ± 0.1 | 3.4% |
| **Intromissions** | S-H- | 2.38 ± 1.70 | 3.13 ± 3.61 | 0.50 ± 1.00 | 3.00 ± 4.76 | 0.13 ± 0.25 | 1.13 ± 1.65 | -52.5% |
| | S+H- | 7.50 ± 9.26 | 1.63 ± 2.93 | 1.50 ± 1.78 | 3.88 ± 5.85 | 2.38 ± 3.09 | 2.00 ± 3.37 | -73.3% |
| | S-H+ | 4.38 ± 3.04 | 6.00 ± 6.36 | 5.88 ± 6.74 | 10.50 ± 8.26 | 5.50 ± 8.82 | 8.75 ± 7.27 | 99.8% |
| | S+H+ | 6.25 ± 12.50 | 9.25 ± 7.27 | 13.38 ± 4.64 | 17.13 ± 10.88 | 9.63 ± 7.45 | 22.25 ± 10.84 | 256.0% |
| | S-H++ | 23.75 ± 8.96 | 11.88 ± 10.94 | 28.00 ± 11.74 | 16.63 ± 11.88 | 10.13 ± 8.80 | 18.00 ± 13.88 | -24.2% |
| **% Time with Male** | S-H- | 18.94 ± 19.58 | 20.09 ± 11.29 | 13.09 ± 19.24 | 18.65 ± 16.45 | 22.65 ± 21.79 | 18.85 ± 20.22 | -0.5% |
| | S+H- | 40.75 ± 13.18 | 25.73 ± 14.01 | 26.42 ± 20.94 | 22.28 ± 16.55 | 27.61 ± 25.83 | 25.86 ± 25.85 | -36.5% |
| | S-H+ | 17.53 ± 14.35 | 21.45 ± 12.28 | 17.33 ± 11.58 | 12.41 ± 10.77 | 22.14 ± 17.16 | 12.22 ± 7.95 | -30.3% |
| | S+H+ | 30.02 ± 11.04 | 18.99 ± 4.67 | 21.99 ± 11.97 | 28.71 ± 13.57 | 27.35 ± 12.56 | 31.69 ± 9.98 | 5.6% |
| | S-H++ | 25.26 ± 6.70 | 26.26 ± 15.23 | 35.01 ± 18.61 | 17.39 ± 17.21 | 21.21 ± 16.91 | 27.27 ± 11.08 | 8.0% |
| **% Time in Doorway** | S-H- | 10.91 ± 7.54 | 16.86 ± 3.57 | 18.60 ± 14.28 | 22.62 ± 11.98 | 19.38 ± 13.43 | 19.87 ± 9.05 | 82.1% |
| | S+H- | 14.18 ± 1.92 | 16.72 ± 6.96 | 22.03 ± 10.88 | 28.73 ± 16.31 | 25.95 ± 12.63 | 21.65 ± 12.09 | 52.7% |
| | S-H+ | 19.95 ± 15.91 | 24.2 ± 12.75 | 26.45 ± 19.49 | 33.6 ± 18.97 | 27.51 ± 17.06 | 21.95 ± 16.71 | 10.0% |
| | S+H+ | 18.78 ± 12.03 | 25.49 ± 12.21 | 23.80 ± 14.72 | 25.66 ± 10.02 | 29.89 ± 17.98 | 29.15 ± 7.79 | 55.2% |
| | S-H++ | 19.23 ± 8.22 | 17.33 ± 10.2 | 15.53 ± 6.64 | 22.97 ± 11.87 | 25.72 ± 17.34 | 24.93 ± 17.21 | 29.6% |
| **% Time Alone** | S-H- | 70.15 ± 24.91 | 63.05 ± 7.74 | 68.31 ± 26.81 | 58.73 ± 27.22 | 57.96 ± 35.03 | 61.28 ± 28.52 | -12.6% |
| | S+H- | 45.07 ± 14.47 | 57.54 ± 16.48 | 51.56 ± 21.23 | 48.99 ± 18.68 | 46.44 ± 29.42 | 52.48 ± 34.62 | 16.4% |
| | S-H+ | 62.52 ± 28.45 | 54.35 ± 24.92 | 56.22 ± 29.41 | 53.99 ± 28.77 | 50.35 ± 33.78 | 65.83 ± 22.53 | 5.3% |
| | S+H+ | 51.20 ± 15.31 | 55.52 ± 12.82 | 54.20 ± 22.77 | 45.63 ± 21.85 | 42.76 ± 24.62 | 39.15 ± 12.09 | -23.5% |
| | S-H++ | 55.51 ± 8.98 | 56.41 ± 22.67 | 49.45 ± 15.52 | 59.64 ± 23.40 | 53.07 ± 32.29 | 47.80 ± 26.25 | -13.9% |
